# Hybrid immunity improves B cell frequency, antibody potency and breadth against SARS-CoV-2 and variants of concern

**DOI:** 10.1101/2021.08.12.456077

**Authors:** Emanuele Andreano, Ida Paciello, Giulia Piccini, Noemi Manganaro, Piero Pileri, Inesa Hyseni, Margherita Leonardi, Elisa Pantano, Valentina Abbiento, Linda Benincasa, Ginevra Giglioli, Concetta De Santi, Massimiliano Fabbiani, Ilaria Rancan, Mario Tumbarello, Francesca Montagnani, Claudia Sala, Emanuele Montomoli, Rino Rappuoli

## Abstract

To understand the nature of the antibody response to SARS-CoV-2 vaccination, we analyzed at single cell level the B cell responses of five naïve and five convalescent people immunized with the BNT162b2 mRNA vaccine. Convalescents had higher frequency of spike protein specific memory B cells and by cell sorting delivered 3,532 B cells, compared with 2,352 from naïve people. Of these, 944 from naïve and 2,299 from convalescents produced monoclonal antibodies against the spike protein and 411 of them neutralized the original Wuhan SARS-CoV-2 virus. More than 75% of the monoclonal antibodies from naïve people lost their neutralization activity against the B.1.351 (beta) and B.1.1.248 (gamma) variants while this happened only for 61% of those from convalescents. The overall loss of neutralization was lower for the B.1.1.7 (alpha) and B.1.617.2 (delta) variants, however it was always significantly higher in those of naïve people. In part this was due to the IGHV2-5;IGHJ4-1 germline, which was found only in convalescents and generated potent and broadly neutralizing antibodies. Overall, vaccination of seropositive people increases the frequency of B cells encoding antibodies with high potency and that are not susceptible to escape by any of the four variants of concern. Our data suggest that people that are seropositive following infection or primary vaccination will produce antibodies with increased potency and breadth and will be able to better control SARS-CoV-2 emerging variants.

## INTRODUCTION

Twenty months after the beginning of the COVID-19 pandemic, with 200 million people infected, 4.2 million deaths, and 3.9 billion vaccines doses administered, the world is still struggling to control the virus. In most developed countries vaccines have vastly reduced severe diseases, hospitalization and deaths, but they have not been able to control the infections which are fueled by new and more infectious variants. A large number of studies so far have shown that protection from infection is linked to the production of neutralizing antibodies against the spike protein (S protein) of the virus^1–4^. This is a metastable, trimeric class 1 fusion glycoprotein, composed by the S1 and S2 subunits, and mediates virus entry changing from a prefusion to postfusion conformation after binding to the human angiotensin-converting enzyme 2 (ACE2) receptor and heparan sulfates on the host cells^5^. Potent neutralizing antibodies recognize the S1 subunit of each monomer which includes the receptor binding domain (RBD) and N-terminal domain (NTD) immunodominant sites^6^. The large majority of neutralizing antibodies bind the receptor binding motif (RBM), within the RBD, and a smaller fraction target the NTD^3,7^. Neutralizing antibodies against the S2 subunit have been described, however they have very low potency^3,8^. Neutralizing antibodies generated after infection derive in large part from germline IGHV3-53 and the closely related IGHV3-66 with very few somatic mutations^9,10^. Starting from the summer of 2020, the virus started to generate mutations that allowed the virus to evade neutralizing antibodies, to become more infectious, or both. Some of the mutant viruses completely replaced the original Wuhan SARS-CoV-2. The most successful variant viruses are the B.1.1.7 (alpha), B.1.351 (beta), B.1.1.248 (gamma) and B.1.617.2 (delta) which have been named Variants of Concern (VoCs) by the World Health Organization^11^. The delta variant is presently spreading across the globe and causing big concern also in fully vaccinated populations. It is therefore imperative to understand the molecular dynamics of the immune response to vaccination in order to design better vaccines and vaccination policies. Several investigators have shown that vaccination of convalescent people can yield neutralizing antibodies which can be up to a thousand-fold higher than those induced by infection or vaccination, suggesting that one way of controlling the pandemic may be the induction of a hybrid immunity-like response using a third booster dose^12–16^. Here we compared at single cell level the nature of the neutralizing antibody response against the original Wuhan virus and the VoCs in naïve and convalescent subjects immunized with the BNT162b2 mRNA vaccine. Naïve subjects were seronegative before vaccination while convalescent donors were already seropositive before vaccination. They will be named seronegatives and seropositives respectively in this work. Our data suggest that immunization of people already seropositive to the virus, increases the frequency, potency and breadth of neutralizing antibodies and may help to control emerging variants.

## RESULTS

### High levels of S protein specific memory B cells and plasma activity in seropositive vaccinees

In this study we enrolled 10 donors vaccinated with the BNT162b2 mRNA vaccine, 5 of them were healthy people naïve to SARS-CoV-2 infection at vaccination (seronegative) and other 5 had recovered from SARS-CoV-2 infection before vaccination (seropositive). Subject details are summarized in **Table S1**. We initially analyzed the B cell population frequencies between our groups. Seropositives showed a 2.46-fold increase in S protein specific CD19^+^CD27^+^IgD^-^IgM^- +^ MBCs compared to seronegatives and an overall 10% higher level of CD19^+^CD27^+^IgD^-^IgM^-^MBCs (**Figure S1A-B**). On the other hand, seronegatives showed a 2.3-fold higher frequency of CD19^+^CD27^+^IgD^-^IgM^+^ MBCs compared to seropositives. No differences were found in the levels of CD19^+^CD27^+^IgD^-^IgM^+^ S protein^+^ MBCs between the two groups assessed in this study (**Figure S1A-B**). Following the MBC analyses, we characterized the polyclonal response of these donors by testing their binding response to the S protein trimer, RBD, NTD and S2-domain, and subsequently by testing their neutralization activity against the original Wuhan SARS-CoV-2 virus (**Figure S2**). Plasma from seropositives showed a higher binding activity to the S protein and all tested domains compared to seronegatives (**Figure S2A – D**). In addition, seropositives showed a 10-fold higher neutralization activity against the original Wuhan SARS-CoV-2 virus compared to seronegatives (**Figure S2E – F**).

### Frequency of neutralizing antibodies against the Wuhan virus and variants of concern

To better characterize the B cell immune response in seronegative and seropositive donors, we single cell sorted antigen-specific memory B cell (MBCs) using as bait the stabilized Wuhan SARS-CoV-2 S protein antigen which was encoded by the mRNA vaccine. The single cell sorting strategy was performed as previously described^3^. Briefly, MBCs prefusion S protein trimer-specific (S protein^+^), class-switched MBCs (CD19^+^CD27^+^IgD^-^IgM^-^) were single-cell sorted and then incubated for two weeks to naturally produce and release mAbs into the supernatant. A total of 2,352 and 3,532 S protein^+^ MBCs were sorted from seronegative and seropositive vaccinees respectively (**Table S2**). Of these 944 (40.1%) and 2,299 (65.1%) respectively, released in the supernatant monoclonal antibodies (mAbs) recognizing the S protein prefusion trimer in ELISA (**Figure 1A**; **Table S2**). These mAbs were then tested in a cytopathic effect-based microneutralization assay (CPE-MN) with the original Wuhan live SARS-CoV-2 virus at a single point dilution (1:5) to identify SARS-CoV-2 neutralizing human monoclonal antibodies (nAbs). This first screening identified a total of 411 nAbs, of which 71 derived from seronegatives and 340 were from seropositives(**Figure 1B**; **Table S2**). Overall, the fraction of S protein-specific B cells producing nAbs were 7.5% for seronegatives and 14.8% for seropositives. Following this first screening, all nAbs able to neutralize the Wuhan SARS-CoV-2 virus were tested by CPE-MN against major variants of concern (VoCs) including the B.1.1.7 (alpha), B.1.351 (beta) and B.1.1.248 (gamma) to understand the breadth of neutralization of nAbs elicited by the BNT162b2 mRNA vaccine. At the time of this assessment the B.1.617.2 (delta) variant was not yet spread globally and therefore it was not available for screening. Seropositives had an overall higher percentage of nAbs neutralizing the VoCs compared to seronegatives. The average frequency of nAbs from seropositives neutralizing the alpha, beta and gamma variants was 80.6 (n=274), 39.4 (n=134) and 62.0% (n=211) respectively, compared to 70.4 (n=50), 22.5 (n=16) and 43.6% (n=31) respectively in seronegatives (**Figure 1C**; **Table S2**).

**Figure 1.**
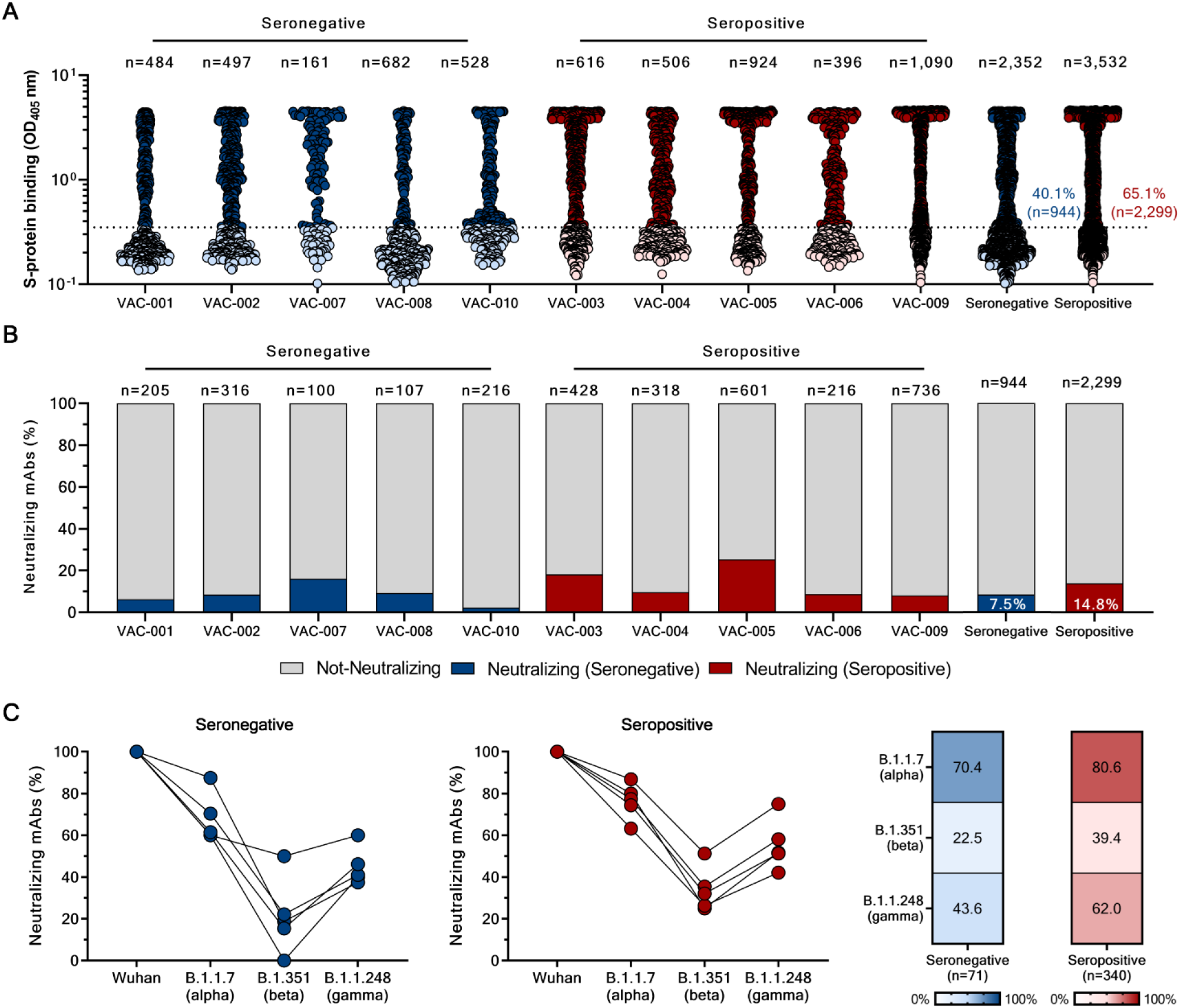
Identification of cross-neutralizing SARS-CoV-2 S protein-specific nAbs. (A) The graph shows supernatants tested for binding to the Wuhan SARS-CoV-2 S protein antigen stabilized in its prefusion conformation. Threshold of positivity has been set as two times the value of the blank (dotted line). Dark blue and red dots represent mAbs that bind to the S protein for seronegative and seropositive vaccinees respectively. Light blue and red dots represent mAbs that do not bind the S protein for seronegative and seropositive vaccinees. (B) The bar graph shows the percentage of not-neutralizing (gray), neutralizing mAbs from seronegatives (dark blue), and neutralizing mAbs for seropositives (dark red). The total number (n) of antibodies tested per individual is shown on top of each bar (C) Graphs show the fold change percentage of nAbs in seronegatives and seropositives against the alpha, beta and gamma VoCs compared to the original Wuhan SARS-CoV-2 virus. The heatmaps show the overall percentage of Wuhan SARS-CoV-2 nAbs able to neutralize tested VoCs.

### High potency and breadth of neutralization in seropositive COVID-19 vaccinees

To better characterize and understand the potency and breadth of coverage of all Wuhan SARS-CoV-2 nAbs, we aimed to express as immunoglobulin G1 (IgG1) all the 411 nAbs previously identified. We were able to recover and express 276 antibodies for further characterization, 224 (89.8%) from seropositives and 52 (10.2%) from seronegatives. Initially, antibodies were tested for binding against the RBD, NTD and the S2-domain of the original Wuhan SARS-CoV-2 S protein. Overall, no major differences were observed in nAbs that recognized the RBD and NTD with 71.2% (n=37) and 79.5% (n=178) nAbs binding the RBD, while 17.3% (n=9) and 16.5% (n=37) nAbs binding the NTD for seronegatives and seropositives respectively (**Figure S3**). None of the tested nAbs targeted the S2 domain. The biggest difference between groups was seen for nAbs able to bind the S protein only in its trimeric conformation i.e. not able to bind single domains. This class of nAbs was almost 3-fold higher in seronegatives compared to seropositives (**Figure S3**). nAbs were then tested by CPE-MN in serial dilution to evaluate their 100% inhibitory concentration (IC_100_) against the Wuhan SARS-CoV-2 virus and the VoCs. At this stage of the study, the B.1.617.2 (delta) spread globally, and we were able to obtain the live virus for our experiments. Overall, nAbs isolated from seropositive vaccinees had a significantly higher potency than those isolated from seronegatives. The IC_100_ geometric mean (GM-IC_100_) in seropositives was 2.87, 2.17-, 1.17-, 1.43-, and 1.92-fold lower than in seronegatives for the Wuhan virus, the alpha, beta, gamma and delta VoCs respectively (**Figure 2**). In addition, a bigger fraction of nAbs from seropositives retained the ability to neutralize the VoCs. Indeed, when nAbs were individually tested against all VoCs, the ability to neutralize the alpha, beta, gamma and delta variants was lost by 14, 61, 61 and 29% of the antibodies from seropositives versus 32, 78, 75 and 46% respectively of those from seronegatives. The overall number of nAbs that lost neutralizing activity against the beta and gamma VoCs was very high, (75-78% in seronegatives and 61% in seropositives), while it was much lower for the alpha and delta variants (32-46% for seronegatives and 14-29% for seropositives) (**Figure 2**). Finally, a major difference between seronegatives and seropositives was found in the class of medium/high potency nAbs (IC_100_ of 11-100 ng/mL and 101–1000 ng/mL) against all variants. Indeed, nAbs in these ranges from seropositives constitute the 71.0%, 62.5%, 23.7%, 22.8%, 53,1% of the whole nAbs repertoire while nAbs from seronegative donors were 48.1%, 38.5%, 17.3%, 17.3%, 34.6% against the Wuhan SARS-CoV-2 virus and alpha, beta, gamma and delta VoCs respectively (**Figure 2**).

**Figure 2.**
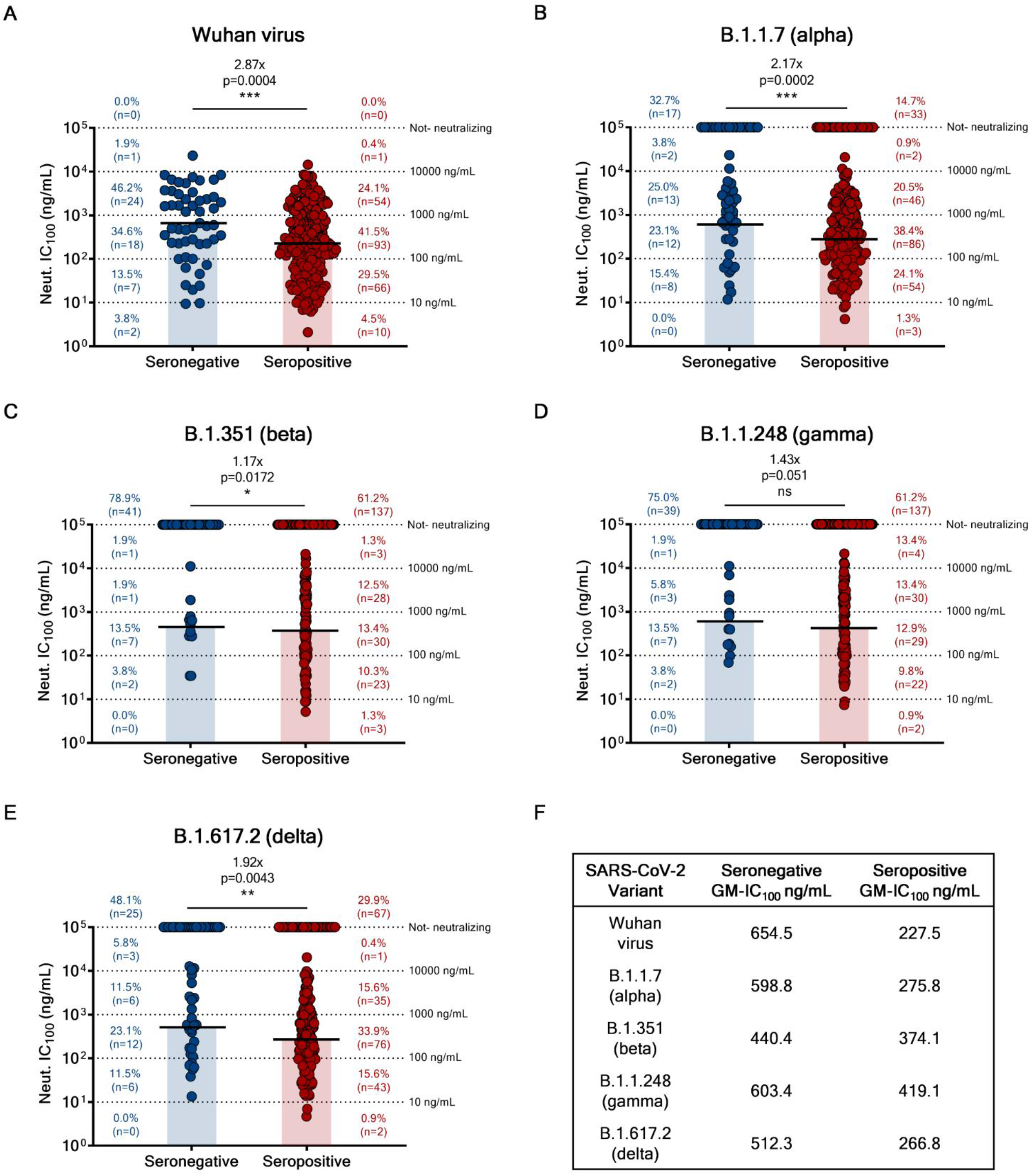
Potency and breadth of neutralization of nAbs against SARS-CoV-2 and VoCs. (A - E) Scatter dot charts show the neutralization potency, reported as IC_100_ (ng/mL), of nAbs tested against the original Wuhan SARS-CoV-2 virus (A) and the B.1.1.7 (B), B.1.351 (C), B.1.1.248 (D) and B.1.617.2 (E) VoCs. The number and percentage of nAbs from seronegatives vs seropositives, fold-change and statistical significance are denoted on each graph. A nonparametric Mann–Whitney t test was used to evaluate statistical significances between groups. Significances are shown as *p < 0.05, **p < 0.01, ***p < 0.001, and ****p < 0.0001. (F) The table shows the IC_100_ geometric mean (GM) of all nAbs pulled together from each group against all SARS-CoV-2 viruses tested.

### Functional gene repertoire of neutralizing antibodies

The analysis of the immunoglobulin G heavy chain variable (IGHV) and joining (IGHJ) gene rearrangements of 58 and 278 sequences recovered from seronegative and seropositive subjects respectively, showed that they use a broad range of germlines and share the most abundant. In particular, both groups predominantly used the IGHV1-69;IGHJ4-1 and IGHV3-53;IGHJ6-1, which were shared by three out five subjects per each group (**Figure 3A**). In addition, the IGHV3-30;IGHJ6-1 and IGHV3-33;IGHJ4-1 germlines, more abundant in seronegative donors, and IGHV1-2;IGHJ6-1, mainly expanded in seropositive vaccinees, were also used with high frequency in both groups. Only the IGHV2-5;IGHJ4-1 germline was seen to be predominantly expanded only in seropositive donors (**Figure 3A**). To better characterize these predominant gene families, we evaluated their neutralization potency and breadth against SARS-CoV-2 and VoCs. In this analyses we could not evaluate IGHV3-33;IGHJ4-1 nAbs, as only three of these antibodies were expressed, but we included the IGHV3-53 closely related family IGHV3-66;IGHJ4-1, as this family was previously described to be mainly involved in SARS-CoV-2 neutralization^9,17^. A large part of nAbs deriving from these predominant germlines had a very broad range of neutralization potency against the original Wuhan SARS-CoV-2 virus with IC_100_ spanning from less than 10 to over 10,000 ng/mL (**Figure 3B – G**). However, many of them lost the ability to neutralize SARS-CoV-2 VoCs. The loss of neutralizing activity occurred for most germlines and it was moderate against the alpha and delta variants, while it was dramatic against the beta and gamma variants (**Figure 3 B – G**). A notable exception was the IGHV2-5;IGHJ4-1 germline, present only in nAbs of seropositive patients, that showed potent antibodies able to equally neutralize all SARS-CoV-2 VoCs (**Figure 3D**). Finally, we evaluated the CDRH3-length and V-gene somatic hyper mutation (SHM) levels for all nAbs retrieved from seronegatives and seropositives and for predominant germlines. Overall, the two groups show a similar average CDRH3-length (15.0 aa and 15.1 aa for seronegatives and seropositives respectively), however seropositives showed almost 2-fold higher V-gene mutation levels compared to seronegatives (**Figure S4**). As for predominant gene derived nAbs, we observed heterogenous CDRH3-length, with the only exception of IGHV3-53;IGHJ6-1 nAbs, and higher V-gene mutation levels in seropositives predominant germlines compared to seronegatives (**Figure S5**).

**Figure 3.**
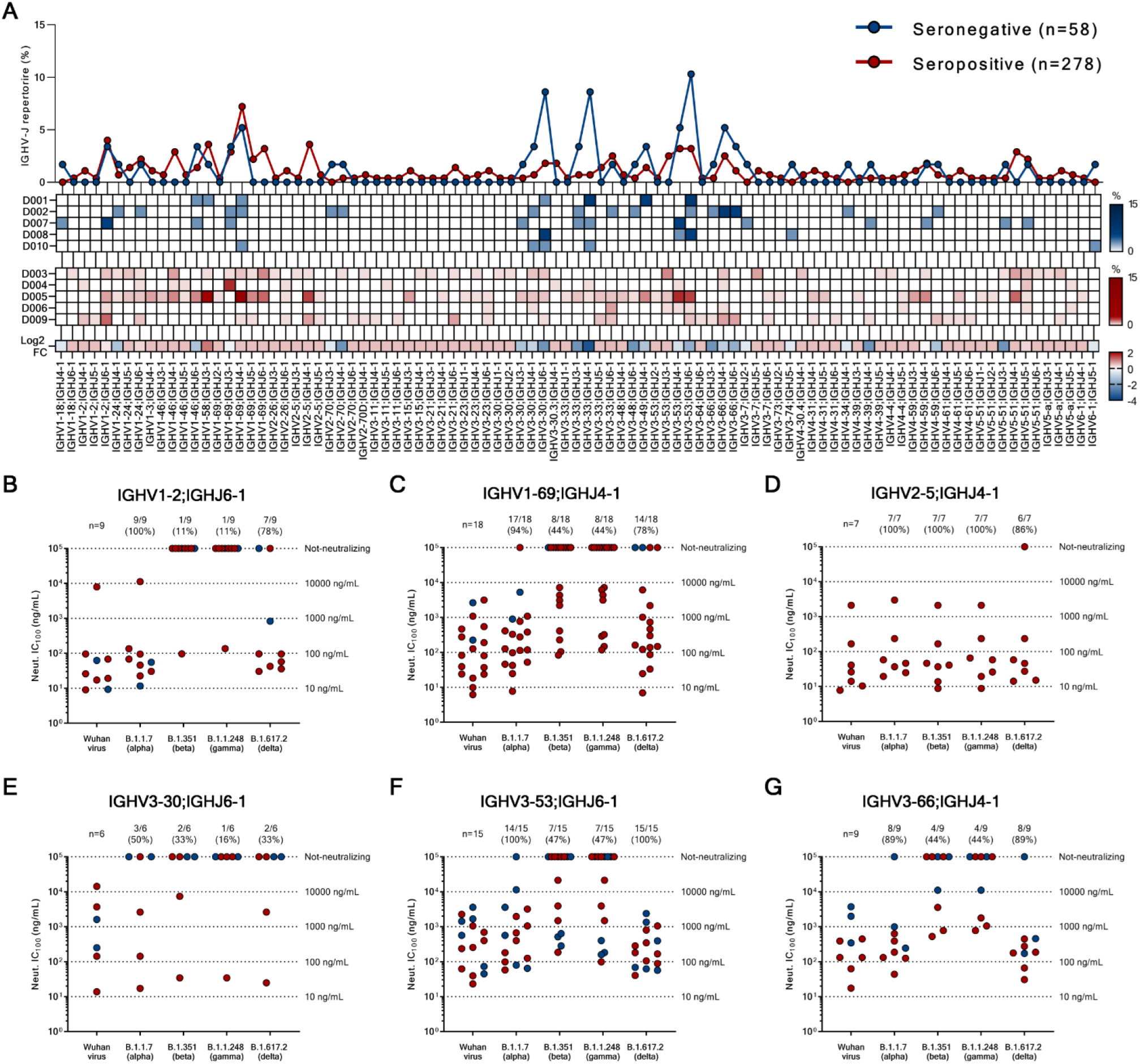
Repertoire analyses and functional characterization of predominant gene derived nAbs. (A) The graph shows the IGHV-J rearrangement frequencies between seronegative and seropositive vaccinees (top panel), the frequency within subjects (middle panels) and the Log2 fold change (FC) between groups (bottom panel). (B – G) Graphs show the neutralization potency (IC_100_) of predominant gene derived nAbs from the IGHV1-2;IGHJ6-1 (B), IGHV1-69;IGHJ4-1 (C), IGHV2-5;IGHJ4-1 (D), IGHV3-30;IGHJ6-1 (E), IGHV3-53;IGHJ6-1 (F) and IGHV3-66;IGHJ4-1 (G) families, against the he original Wuhan SARS-CoV-2 virus and the B.1.1.7, B.1.351, B.1.1.248 and B.1.617.2 VoCs.

### Epitope mapping of neutralizing antibodies

To map the regions of the S protein recognized by the identified nAbs we used a competition assay with four known antibodies: J08, which targets the top loop of the receptor binding motif (RBM)^3^, S309, which binds the RBD but outside of the RMB region^18^, 4A8, that recognized the NTD^19^, and L19, that binds the S2-domain^3^(**Figure S6**). The nAbs identified in this study were pre-incubated with the original Wuhan SARS-CoV-2 S protein and subsequently the four nAbs labeled with different fluorophores were added as single mix. 50% signal reduction for one of the four fluorescently labelled nAbs, was used as threshold for positive competition. The vast majority of nAbs from both seronegative (50.0%; n=26) and seropositive (51.3%; n=115) vaccinees competed with J08 (**Figure 4A; Table S3**). For seronegatives, the second most abundant population was composed by nAbs that did not compete with any of the four fluorescently labelled nAbs (25.0%; n=13) followed by nAbs targeting the NTD (17.3%; n=9). As for seropositives, the second most abundant population was composed by nAbs that competed with S309 (21.4%; n=48) followed by nAbs competing with 4A8 (15.6%; n=35) and not-competing nAbs (11.6%; n=26). None of our nAbs did compete with the S2 targeting antibody L19 (**Figure 4A; Table S3**). nAbs competing with J08, which are likely to bind the RBM, derived from several germlines, including the predominant IGHV3-53;IGHJ6-1 (10.6%; n=14), IGHV1-69;IGHJ4-1 (8.3%; n=11) and IGHV1-2;IGHJ6-1 (6.8%; n=9) (**Figure 4B**). In contrast, those competing with S309 derived mostly from germline IGHV2-5;IGHJ4-1 (13.7%; n=7) which were isolated exclusively from seropositive vaccinees (**Figure 4C**). As for NTD-directed nAbs the non-predominant gene family IGHV1-24;IGHJ6-1 was the most abundant confirming what was reported in previous studies (**Figure 4D**)^20^. Finally, for nAbs that did not compete with any of the known antibodies used in our competition assay, the non-predominant gene families IGHV1-69;IGHJ3-1 (9.7%; n=3) and IGHV1-69;IGHJ6-1 (9.7%; n=3) were the most abundant (**Figure 4E**).

**Figure 4.**
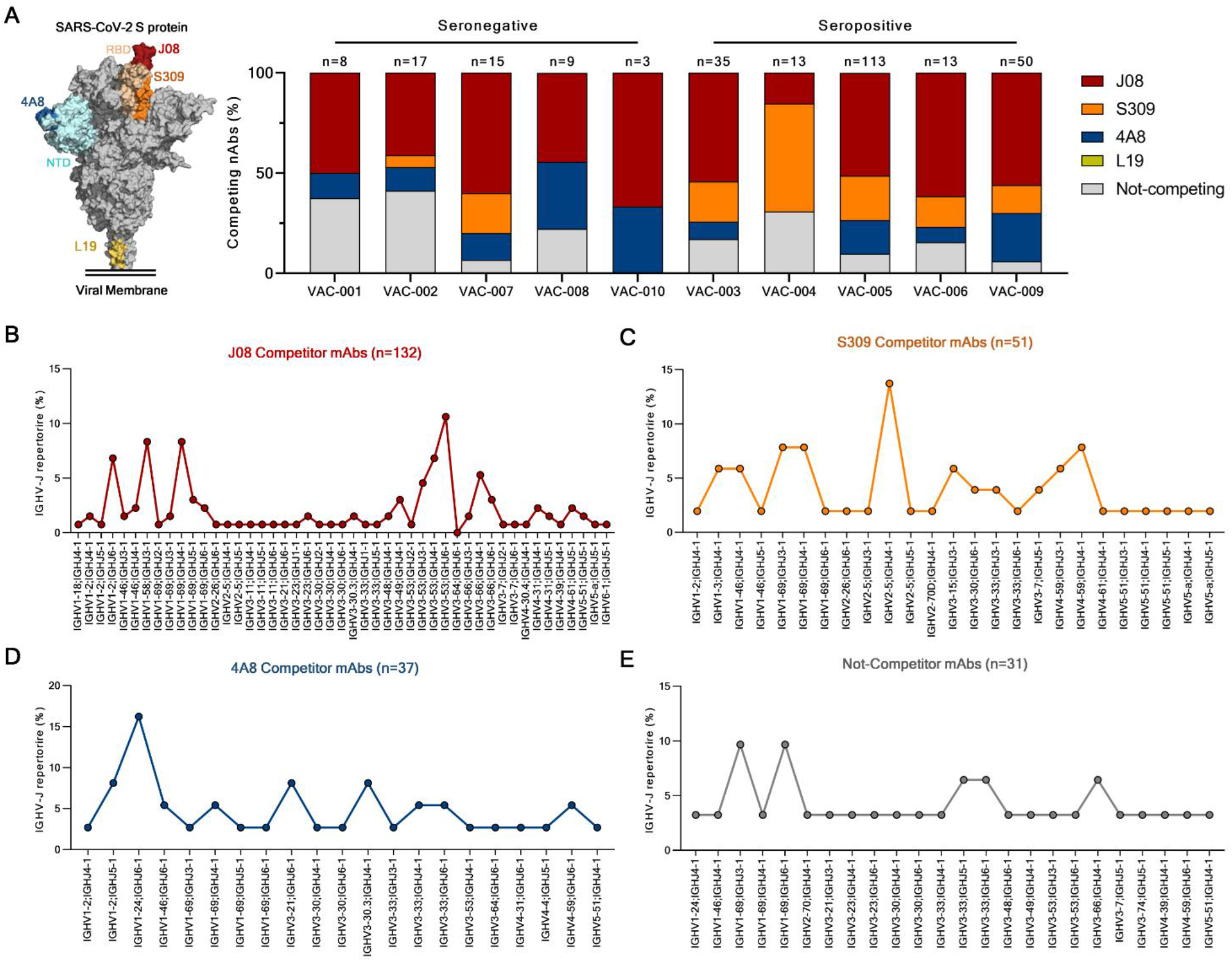
Epitope binning and genetic characterization of competing nAbs. (A) The bar graph shows the percentage (%) of nAbs competing with J08 (dark red), S309 (orange), 4A8 (dark blue) and L19 (gold), or antibodies that did not compete with any of the previous mAbs (gray). A schematic representation of J08, S309, 4A8 and L19 epitopes on the S protein surface is shown on the left side of the panel. (B – D) Graphs show the IGHV-J rearrangement percentage for nAbs that competed against J08 (B), S309 (C), 4A8 (D), or that did not compete with any of these mAbs (E). The total number (n) of competing nAbs per group is shown on top of each graph.

## DISCUSSION

Our study analyzed for the first time at single cell level the repertoire of B cells producing neutralizing antibodies following vaccination of naïve and previously infected people. The most important conclusion from this work is that vaccinating people that are already seropositive to the virus, because of previous SARS-CoV-2 infection, induces neutralizing antibodies which are more potent and less susceptible to escape from SARS-CoV-2 variants. This conclusion is not surprising since several papers have already reported that vaccination of convalescent people induces a hybrid immunity with titers of neutralizing antibodies up to 50-fold higher than those induced by vaccination of naïve people^15,16^. The novelty of our work lies in the molecular analysis of this response which shows that seropositive subjects respond to vaccination with more B cells producing antibodies which show higher neutralization potency and are not susceptible to escape variants. In part this is due to potent antibodies derived from the IGHV2-5;IGHJ4-1 germline which was only found in seropositive people. The absence of this germline in naïve vaccinees is also confirmed by previous studies^16^. One limitation of our study is that we did not include people that received a third booster dose of vaccine, as at the time that this work was being performed no policies for a third booster dose were implemented. In spite of this limitation, we believe that our conclusions are likely to be extendable to people that are seropositives following primary vaccination. Indeed, the S protein produced following vaccination with mRNA, viral vectors or following infection is produced in all cases by the cell of the host and it is likely to be presented to the immune system in a similar manner and generate a similar antibody response. This is confirmed by the fact that neutralizing antibodies following infection and vaccination derive mostly from the same immunodominant germlines, i.e. IGHV3-53, 3-30 and 3-66^9,10,16,17^. Our analysis suggests that a booster dose of vaccine will increase the frequency of memory B cells producing potent neutralizing antibodies not susceptible to escape variants and allow better control of this pandemic. Our work also shows that more than three quarters of antibodies neutralizing the Wuhan virus do not neutralize at all the beta and gamma variants, while the fraction of antibodies not neutralizing the alpha and delta variants is much smaller and in all cases below 50%. This suggest that the beta and gamma variants were originally selected to escape natural immunity, while the alpha and gamma were selected mostly for their increased infectivity and ability to spread in the population^21,22^. The massive escape from predominant germlines, such as IGHV3-53, 3-66, 3-30 and 1-69, and the presence of antibodies deriving from germline IGHV2-5 that are completely insensitive to the existing variants, suggest that the design of vaccines that preferentially promote or avoid the expansion of selected germlines can generate broad protection against SARS-CoV-2 variants. Germline-targeting vaccination, which has been pioneered in the HIV field^23,24^, may be a promising strategy to fight the COVID-19 pandemic.

## MATERIALS & METHODS

### Enrollment of COVID-19 vaccinees and human sample collection

This work results from a collaboration with the Azienda Ospedaliera Universitaria Senese, Siena (IT) that provided samples from COVID-19 vaccinated donors, of both sexes, who gave their written consent. The study was approved by local ethics committees (Parere 17065 in Siena) and conducted according to good clinical practice in accordance with the declaration of Helsinki (European Council 2001, US Code of Federal Regulations, ICH 1997). This study was unblinded and not randomized. No statistical methods were used to predetermine sample size.

### Single cell sorting of SARS-CoV-2 S protein^+^ memory B cells from COVID-19 vaccinees

Peripheral blood mononuclear cells (PBMCs) and single cell sorting strategy were performed as previously described^3^. Briefly, PBMC were isolated from heparin-treated whole blood by density gradient centrifugation (Ficoll-Paque™ PREMIUM, Sigma-Aldrich). After separation, PBMC were stained with Live/Dead Fixable Aqua (Invitrogen; Thermo Scientific) diluted 1:500 at room temperature RT. After 20 min incubation cells were washed with PBS and unspecific bindings were saturated with 20% normal rabbit serum (Life technologies). Following 20 min incubation at 4°C cells were washed with PBS and stained with SARS-CoV-2 S-protein labeled with Strep-Tactin®XT DY-488 (iba-lifesciences cat# 2-1562-050) for 30 min at 4°C. After incubation the following staining mix was used CD19 V421 (BD cat# 562440), IgM PerCP-Cy5.5 (BD cat# 561285), CD27 PE (BD cat# 340425), IgD-A700 (BD cat# 561302), CD3 PE-Cy7 (BioLegend cat# 300420), CD14 PE-Cy7 (BioLegend cat# 301814), CD56 PE-Cy7 (BioLegend cat# 318318) and cells were incubated at 4°C for additional 30 min. Stained MBCs were single cell-sorted with a BD FACS Aria III (BD Biosciences) into 384-well plates containing 3T3-CD40L feeder cells and were incubated with IL-2 and IL-21 for 14 days as previously described^25^.

### ELISA assay with SARS-CoV-2 S protein prefusion trimer

mAbs and plasma binding specificity against the S-protein trimer was detected by ELISA as previously described^3^. Briefly, 384-well plates (microplate clear, Greiner Bio-one) were coated with 3 µg/mL of streptavidin (Thermo Fisher) diluted in carbonate-bicarbonate buffer (E107, Bethyl laboratories) and incubated at RT overnight. The next day, plates were incubated 1 h at RT with 3µg/mL of SARS-CoV-2 S protein diluted in PBS. Plates were then saturated with 50 µL/well of blocking buffer (phosphate-buffered saline, 1% BSA) for 1 h at 37°C. After blocking, 25 µL/well of mAbs diluted 1:5 in sample buffer (phosphate-buffered saline, 1% BSA, 0.05% Tween-20) were added to the plates and were incubated at 37°C. Plasma samples derived from vaccinees were tested (starting dilution 1:10; step dilution 1:2 in sample buffer) in a final volume of 25 µL/well and were incubated at 37°C. After 1 h of incubation, 25 µl/well of alkaline phosphatase-conjugated goat antihuman IgG and IgA (Southern Biotech) diluted 1:2000 in sample buffer were added. Finally, S protein binding was detected using 25 µL/well of PNPP (p-nitrophenyl phosphate;Thermo Fisher) and the reaction was measured at a wavelength of 405 nm by the Varioskan Lux Reader (Thermo Fisher Scientific). After each incubation step, plates were washed three times with 100µl/well of washing buffer (phosphate-buffered saline, 0.05% Tween-20). Sample buffer was used as a blank and the threshold for sample positivity was set at 2-fold the optical density (OD) of the blank.

### ELISA assay with RBD, NTD and S2 subunits

mAbs identification and plasma screening of vaccinees against RBD, NTD or S2 SARS-CoV-2 protein were performed by ELISA. Briefly, 3 µg/mL of RBD, NTD or S2 SARS-CoV-2 protein diluted in carbonate-bicarbonate buffer (E107, Bethyl laboratories) were coated in 384-well plates (microplate clear, Greiner Bio-one). After overnight incubation at 4°C, plates were washed 3 times with washing buffer (phosphate-buffered saline, 0.05% Tween-20) and blocked with 50 µL/well of blocking buffer (phosphate-buffered saline, 1% BSA) for 1h at 37°C. After washing, plates were incubated 1 h at 37 °C with mAbs diluted 1:5 in samples buffer (phosphate-buffered saline, 1% BSA, 0.05% Tween-20) or with plasma at a starting dilution 1:10 and step diluted 1:2 in sample buffer. Wells with no sample added were consider blank controls. Anti-Human IgG −Peroxidase antibody (Fab specific) produced in goat (Sigma) diluted 1:45000 in sample buffer was then added and samples were incubated for 1 h at 37°C. Plates were then washed, incubated with TMB substrate (Sigma) for 15 min before adding the stop solution (H_2_SO_4_ 0.2M). The OD values were identified using the Varioskan Lux Reader (Thermo Fisher Scientific) at 450 nm. Each condition was tested in triplicate and samples tested were considered positive if OD value was 2-fold the blank.

### Flow cytometry-based competition assay

To classify mAbs candidates on the basis of their interaction with Spike epitopes, we performed a flow cytometry-based competition assay. In detail, magnetic beads (Dynabeads His-Tag, Invitrogen) were coated with histidine tagged S protein according to the manufacturers’ instructions. Then, 20 µg/ml of coated S-protein-beads were pre-incubated with unlabeled nAbs candidates diluted 1:2 in PBS for 40 minutes at RT. After incubation, the mix Beads-antibodies was washed with 100 µL of PBS-BSA 1%. Then, to analyze epitope competition, mAbs able to bind RBD (J08, S309), NTD (4A8) or S2 (L19) domain of the S-protein were labeled with 4 different fluorophores (Alexa Fluor 647, 488, 594 and 405) using Alexa Fluor NHS Ester kit (Thermo Scientific), were mixed and incubated with S-protein-beads. Following 40 minutes of incubation at RT, the mix Beads-antibodies was washed with PBS, resuspended in 150 µL of PBS-BSA 1% and analyzed using BD LSR II flow cytometer (Becton Dickinson). Beads with or without S-protein incubated with labeled antibodies mix were used as positive and negative control respectively. Analysis was performed using FlowJo (version 10).

### SARS-CoV-2 authentic viruses neutralization assay

All SARS-CoV-2 authentic virus neutralization assays were performed in the biosafety level 3 (BSL3) laboratories at Toscana Life Sciences in Siena (Italy) and Vismederi Srl, Siena (Italy). BSL3 laboratories are approved by a Certified Biosafety Professional and are inspected every year by local authorities. To evaluate the neutralization activity of identified nAbs against SARS-CoV-2 and all VoCs and evaluate the breadth of neutralization of this antibody is a cytopathic effect-based microneutralization assay (CPE-MN) was performed^3^. Briefly, the CPE-based neutralization assay sees the co-incubation of J08 with a SARS-CoV-2 viral solution containing 100 TCID_50_ of virus and after 1 hour incubation at 37°C, 5% CO_2_. The mixture was then added to the wells of a 96-well plate containing a sub-confluent Vero E6 cell monolayer. Plates were incubated for 3-4 days at 37°C in a humidified environment with 5% CO_2_, then examined for CPE by means of an inverted optical microscope by two independent operators. All nAbs were tested a starting dilution of 1:5 and the IC_100_ evaluated based on their initial concentration while plasma samples were tested starting from a 1:10 dilution. Both nAbs and plasma samples were then diluted step 1:2. Technical duplicates were performed for both nAbs and plasma samples. In each plate positive and negative control were used as previously described^3^.

### SARS-CoV-2 virus variants CPE-MN neutralization assay

The SARS-CoV-2 viruses used to perform the CPE-MN neutralization assay were the original Wuhan SARS-CoV-2 virus (SARS-CoV-2/INMI1-Isolate/2020/Italy: MT066156), SARS-CoV-2 B.1.1.7 (INMI GISAID accession number: EPI_ISL_736997), SARS-CoV-2 B.1.351 (EVAg Cod: 014V-04058), B.1.1.248 (EVAg CoD: 014V-04089) and B.1.617.2 (GISAID ID: EPI_ISL_2029113)^26^.

### Single cell RT-PCR and Ig gene amplification and transcriptionally active PCR expression

The whole process for nAbs heavy and light chain recovery, amplification and transcriptionally active PCR (TAP) expression was performed as previously described^3^. Briefly, 5 µL of cell lysate were mixed with 1 µL of random hexamer primers (50 ng/µL), 1 µL of dNTP-Mix (10 mM), 2 µL 0.1 M DTT, 40 U/µL RNAse OUT, MgCl_2_ (25 mM), 5x FS buffer and Superscript IV reverse transcriptase (Invitrogen) to perform RT-PCR. Reverse transcription (RT) reaction was performed at 42°C/10’, 25°C/10’, 50°C/60’ and 94°/5’. Two rounds of PCR were performed to obtain the heavy (VH) and light (VL) chain amplicons. All PCR reactions were performed in a nuclease-free water (DEPC) in a total volume of 25 µL/well. For PCR I, 4 µL of cDNA were mixed with 10 µM of VH and 10 µM VL primer-mix, 10mM dNTP mix, 0.125 µL of Kapa Long Range Polymerase (Sigma), 1.5 µL MgCl2 and 5 µL of 5x Kapa Long Range Buffer. PCRI reaction was performed at 95°/3’, 5 cycles at 95°C/30’’, 57°C/30’’, 72°C/30’’ and 30 cycles at 95°C/30’’, 60°C/30’’, 72°C/30’’ and a final extension of 72°/2’. Nested PCR II was performed as above starting from 3.5 μL of unpurified PCR I product. PCR II products were purified by Millipore MultiScreen® PCRμ96 plate according to manufacture instructions and eluted in 30 μL of nuclease-free water (DEPC). As for TAP expression, vectors were initially digested using restriction enzymes AgeI, SalI and Xho as previously described and PCR II products ligated by using the Gibson Assembly NEB into 25 ng of respective human Igγ1, Igκ and Igλ expression vectors^27,28^. TAP reaction was performed using 5 μL of Q5 polymerase (NEB), 5 μL of GC Enhancer (NEB), 5 μL of 5X buffer,10 mM dNTPs, 0.125 µL of forward/reverse primers and 3 μL of ligation product, using the following cycles: 98°/2’, 35 cycles 98°/10’’, 61°/20’’, 72°/1’ and 72°/5’.

### Functional repertoire analyses

nAbs VH and VL sequence reads were manually curated and retrieved using CLC sequence viewer (Qiagen). Aberrant sequences were removed from the data set. Analyzed reads were saved in FASTA format and the repertoire analyses was performed using Cloanalyst (http://www.bu.edu/computationalimmunology/research/software/)^29,30^.

### Statistical analysis

Statistical analysis was assessed with GraphPad Prism Version 8.0.2 (GraphPad Software, Inc., San Diego, CA). Nonparametric Mann-Whitney t test was used to evaluate statistical significance between the two groups analyzed in this study. Statistical significance was shown as * for values ≤ 0.05, ** for values ≤ 0.01, *** for values ≤ 0.001, and **** for values ≤ 0.0001.

## Data availability

All data supporting the findings in this study are available within the article or can be obtained from the corresponding author upon request.

## ACKNOWLEDGMENTS

This work was funded by the European Research Council (ERC) advanced grant agreement number 787552 (vAMRes). This publication was supported by funds from the “Centro Regionale Medicina di Precisione” and by all the people who answered the call to fight with us the battle against SARS-CoV-2 with their kind donations on the platform ForFunding (https://www.forfunding.intesasanpaolo.com/DonationPlatform-ISP/nav/progetto/id/3380). This work was funded by COOP ITALIA Soc. Coop. This publication was supported by the European Virus Archive goes Global (EVAg) project, which has received funding from the European Union’s Horizon 2020 research and innovation program under grant agreement No 653316. This publication was supported by the COVID-2020-12371817 project, which has received funding from the Italian Ministry of Health. We would also like to acknowledge Dr. Jason McLellan, for kindly providing the S protein trimer, RBD, NTD and S2 constructs, and Dr. Olivier Schwartz, for providing the B.1.617.2 (delta) SARS-CoV-2 variant. We would like to thank the nurse staff of the operative unit of the department of Medical Sciences, Infectious and Tropical Diseases Unit, Siena University Hospital, Siena, Italy, and all the COVID-19 vaccinated donors for participating to this study.

## AUTHOR CONTRIBUTIONS

EA and RR conceived the project. FM, MF, IR and MT enrolled COVID-19 vaccinees to the study. EA and IP performed PBMC isolation and single cell sorting. IP performed ELISAs and competition assays. IP and NM recovered nAbs VH and VL and expressed antibodies. PP and EA recovered VH and VL sequences and performed the repertoire analyses. EP and VA produced and purified SARS-CoV-2 S protein constructs. EA, GP, IH, ML, LB and GG performed neutralization assays in BSL3 facilities. CDS supported day-by-day laboratory activities and management. EA and RR wrote the manuscript. All authors contributed to the final revision of the manuscript. EA, CS, EM and RR coordinated the project.

## SUPPLEMENTARY TABLES

**Table S1.**
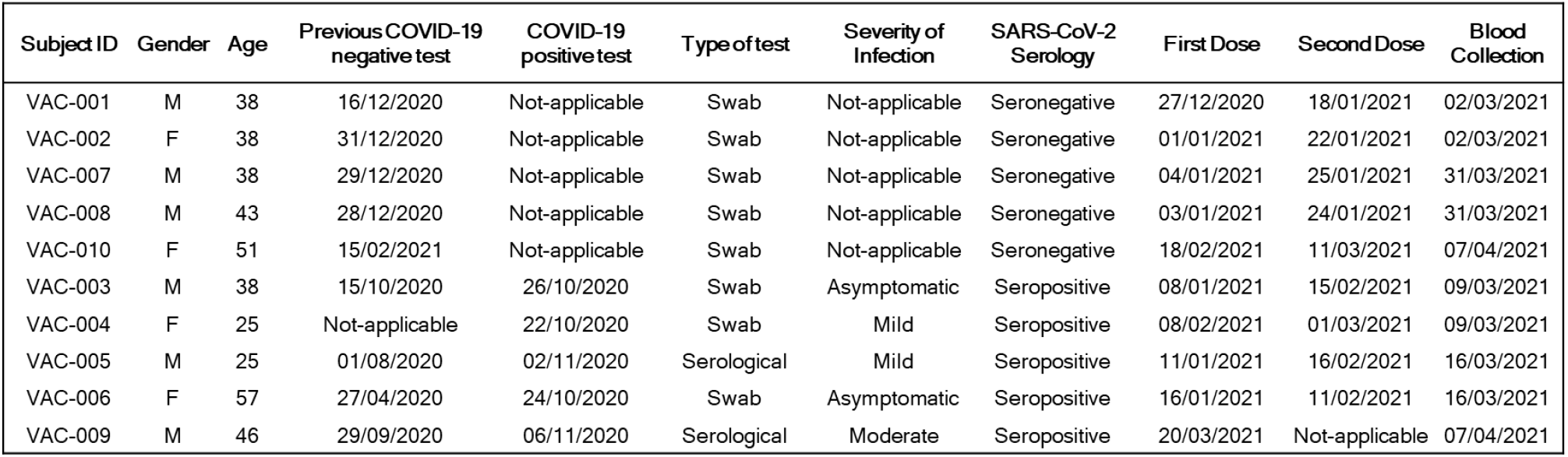
Clinical details of COVID-19 vaccinees. This table summarizes all the clinically relevant information for the 5 seropositive and 5 seronegative vaccinated donors enrolled in this study.

**Table S2.** Summary of COVID-19 vaccinees. This table summarizes the number and percentages of single cell sorted S protein-specific MBCs, number and percentages of S protein binding mAbs, and neutralizing mAbs against the original Wuhan SARS-CoV-2 virus and VoCs. This table includes only data from the initial screening where nAbs were tested at a single point dilution against the original Wuhan virus and the B.1.1.7, B.1.351 and B.1.1.248 VoCs. At this stage the B.1.617.2 (delta) variant was not available.

| Subject | SARS-CoV-2 serology | S-protein+ MBCs Sorted | S-protein+ mAbs (n) | S-protein+ mAbs (%) | Wuhan - Neutralizing antibodies (n) | Wuhan - Neutralizing antibodies (%) | B.1.1.7 - Neutralizing antibodies (n) | B.1.1.7 - Neutralizing antibodies (%) | B.1.351 - Neutralizing antibodies (n) | B.1.351 - Neutralizing antibodies (%) | B.1.1.248 - Neutralizing antibodies (n) | B.1.1.248 - Neutralizing antibodies (%) |
| --- | --- | --- | --- | --- | --- | --- | --- | --- | --- | --- | --- | --- |
| VAC-001 | Seronegative | 484 | 205 | 42.3 | 13 | 6.3 | 8 | 61.5 | 2 | 15.4 | 6 | 46.2 |
| VAC-002 | Seronegative | 497 | 316 | 63.6 | 27 | 8.5 | 19 | 70.4 | 6 | 22.2 | 11 | 40.7 |
| VAC-007 | Seronegative | 161 | 100 | 62.1 | 16 | 16.0 | 14 | 87.5 | 3 | 18.8 | 6 | 37.5 |
| VAC-008 | Seronegative | 682 | 107 | 15.7 | 10 | 9.3 | 6 | 60.0 | 5 | 50.0 | 6 | 60.0 |
| VAC-010 | Seronegative | 528 | 216 | 40.9 | 5 | 2.3 | 3 | 60.0 | 0 | 0.0 | 2 | 40.0 |
| Total (Seronegative) |  | 2,352 | 944 | 40.1 | 71 | 7.5 | 50 | 70.4 | 16 | 22.5 | 31 | 43.6 |
| VAC-003 | Seropositive | 616 | 428 | 69.5 | 78 | 18.2 | 58 | 74.4 | 25 | 32.1 | 40 | 51.3 |
| VAC-004 | Seropositive | 506 | 318 | 62.8 | 31 | 9.7 | 24 | 77.4 | 11 | 35.5 | 18 | 58.1 |
| VAC-005 | Seropositive | 924 | 601 | 65.0 | 152 | 25.3 | 132 | 86.8 | 78 | 51.3 | 114 | 75.0 |
| VAC-006 | Seropositive | 396 | 216 | 55.8 | 19 | 8.8 | 12 | 63.2 | 5 | 26.3 | 8 | 42.1 |
| VAC-009 | Seropositive | 1,090 | 736 | 67.5 | 60 | 8.1 | 48 | 80.0 | 15 | 25.0 | 31 | 51.7 |
| Total (Seropositive) |  | 3,532 | 2,299 | 65.1 | 340 | 14.8 | 274 | 80.6 | 134 | 39.4 | 211 | 62.0 |

**Table S3.** Competition assay. This table summarizes the number and percentages of competing nAbs from seronegative and seropositive vaccinees against J08, S309, 4A8 and L19. Not-competing nAbs are also reported in this table.

| Subject | SARS-CoV-2 serology | Distribution Competition J08 (n) | Distribution Competition J08 (%) | Distribution Competition S309 (n) | Distribution Competition S309 (%) | Distribution Competition 4A8 (n) | Distribution Competition 4A8 (%) | Distribution Competition L19 (n) | Distribution Competition L19 (%) | Distribution Competition Not-competing (n) | Distribution Competition Not-competing (%) |
| --- | --- | --- | --- | --- | --- | --- | --- | --- | --- | --- | --- |
| VAC-001 | Seronegative | 4 | 50.0 | 0 | 0.0 | 1 | 12.5 | 0 | 0.0 | 3 | 37.5 |
| VAC-002 | Seronegative | 7 | 41.2 | 1 | 5.9 | 2 | 11.8 | 0 | 0.0 | 7 | 41.2 |
| VAC-007 | Seronegative | 9 | 60.0 | 3 | 20.0 | 2 | 13.3 | 0 | 0.0 | 1 | 6.7 |
| VAC-008 | Seronegative | 4 | 44.4 | 0 | 0.0 | 3 | 33.3 | 0 | 0.0 | 2 | 22.2 |
| VAC-010 | Seronegative | 2 | 66.7 | 0 | 0.0 | 1 | 33.3 | 0 | 0.0 | 0 | 0.0 |
| Total (Seronegative) |  | 26 | 50.0 | 4 | 7.7 | 9 | 17.3 | 0 | 0.0 | 13 | 25.0 |
| VAC-003 | Seropositive | 19 | 54.3 | 7 | 20.0 | 3 | 8.6 | 0 | 0.0 | 6 | 17.1 |
| VAC-004 | Seropositive | 2 | 15.4 | 7 | 53.8 | 0 | 0.0 | 0 | 0.0 | 4 | 30.8 |
| VAC-005 | Seropositive | 58 | 51.3 | 25 | 22.1 | 19 | 16.8 | 0 | 0.0 | 11 | 9.7 |
| VAC-006 | Seropositive | 8 | 61.5 | 2 | 15.4 | 1 | 7.7 | 0 | 0.0 | 2 | 15.4 |
| VAC-009 | Seropositive | 28 | 56.0 | 7 | 14.0 | 12 | 24.0 | 0 | 0.0 | 3 | 6.0 |
| Total (Seropositive) |  | 115 | 51.3 | 48 | 21.4 | 35 | 15.6 | 0 | 0.0 | 26 | 11.6 |

## SUPPLEMENTARY FIGURES

**Figure S1.**
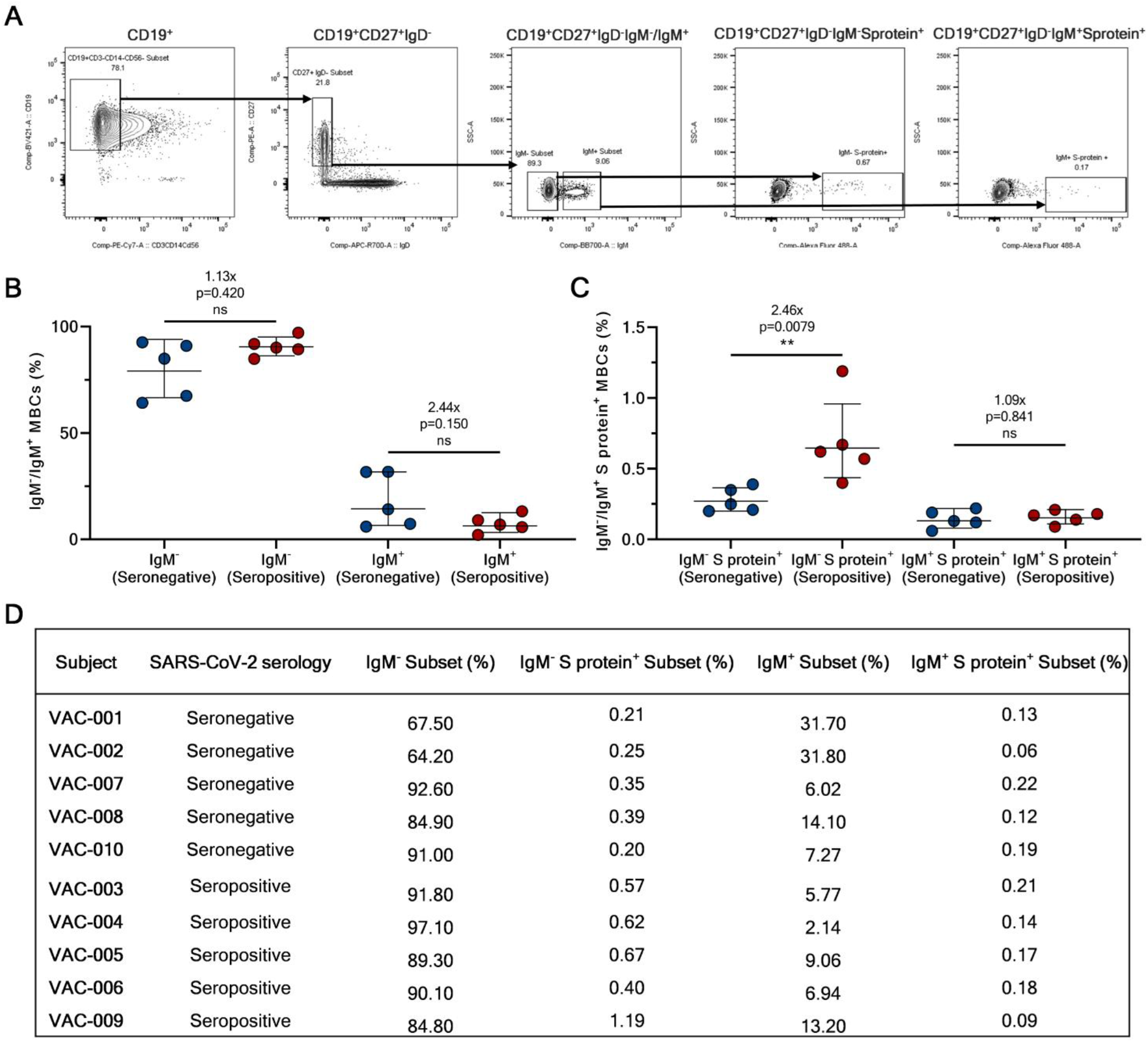
Single cell sorting and memory B cell frequencies. (A) The gating strategy shows from left to right: CD19^+^ B cells; CD19^+^CD27^+^IgD^-^; CD19^+^CD27^+^IgD^-^IgM^-^/IgM^+^; CD19^+^CD27^+^IgD^-^IgM^-^Sprotein^+^; CD19^+^CD27^+^IgD^-^IgM^+^Sprotein^+^. (B) The graph shows the frequency of CD19^+^CD27^+^IgD^-^IgM^-^ and IgM^+^. (C) The graph shows the frequency of CD19^+^CD27^+^IgD^-^IgM^-^ and IgM^+^ able to bind the SARS-CoV-2 S protein trimer (S protein^+^). (D) The table summarizes the frequencies of the cell population above described for all subjects enrolled in our study.

**Figure S2.**
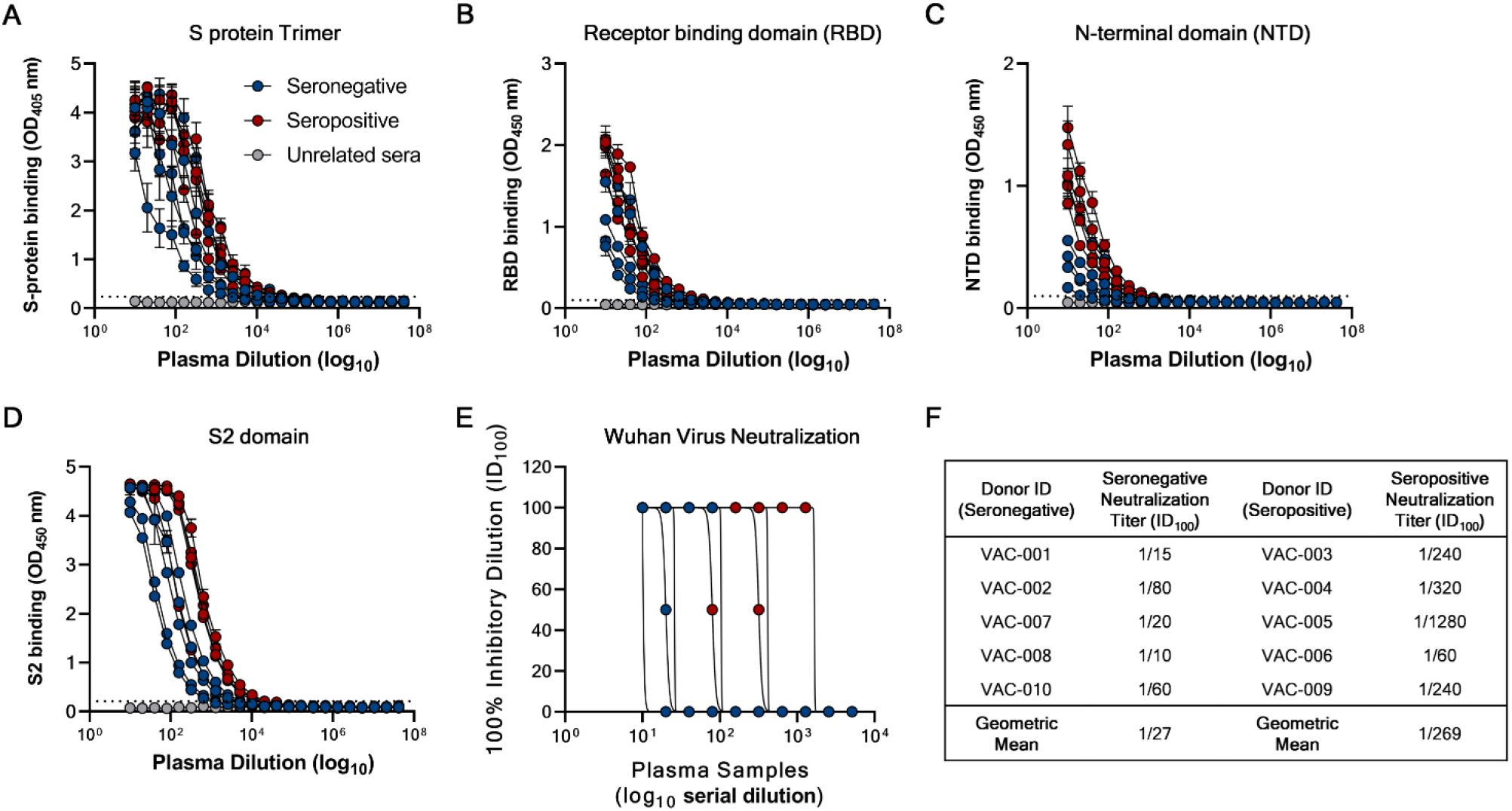
Plasma response of COVID-19 vaccinees. (A - D) Graphs show the ability of plasma samples from seronegative and seropositive vaccinees to bind the S protein trimer (A), RBD (B), NTD (C) and S2 domain (D). (E) The graph shows the neutralizing activity of plasma samples against the original Wuhan SARS-CoV-2 virus. (F) The table summarizes the 100% inhibitory dilution (ID_100_) of each COVID-19 vaccinee and the geometric mean for seronegative and seropositive donors.

**Figure S3.**
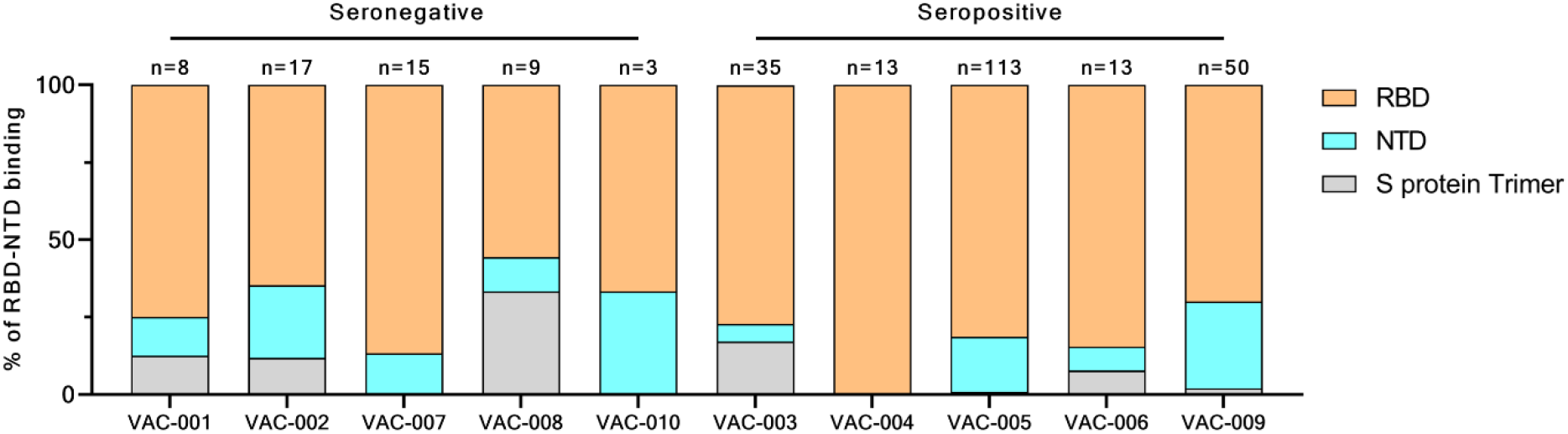
RBD and NTD binding distribution of nAbs. The graph shows the percentage of antibodies that bind specifically the RBD (light orange) or the NTD (cyan) or that did not bind single domains but recognized exclusively the S protein in its trimetric conformation (gray).

**Figure S4.**
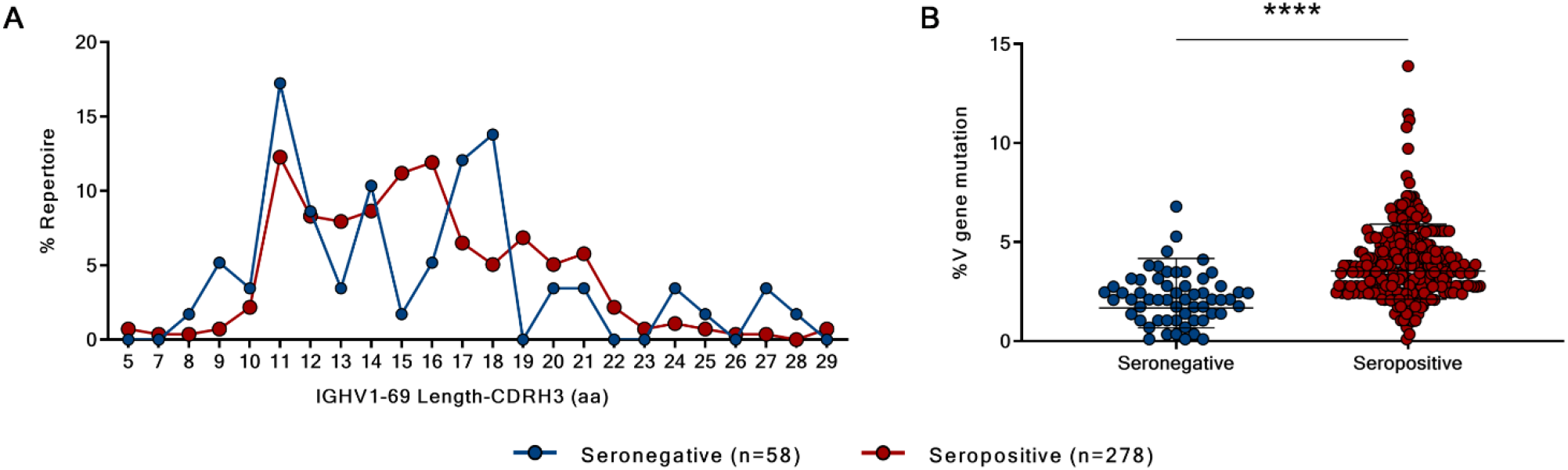
Heavy chain CDR3 length and somatic hypermutation levels in seronegative and seropositive vaccinees. (A) The graph shows the heavy chain CDR3 length represented in amino acids (aa). (B) The graph shows the overall somatic hypermutation level of nAbs isolated from seronegative and seropositive vaccinees.

**Figure S5.**
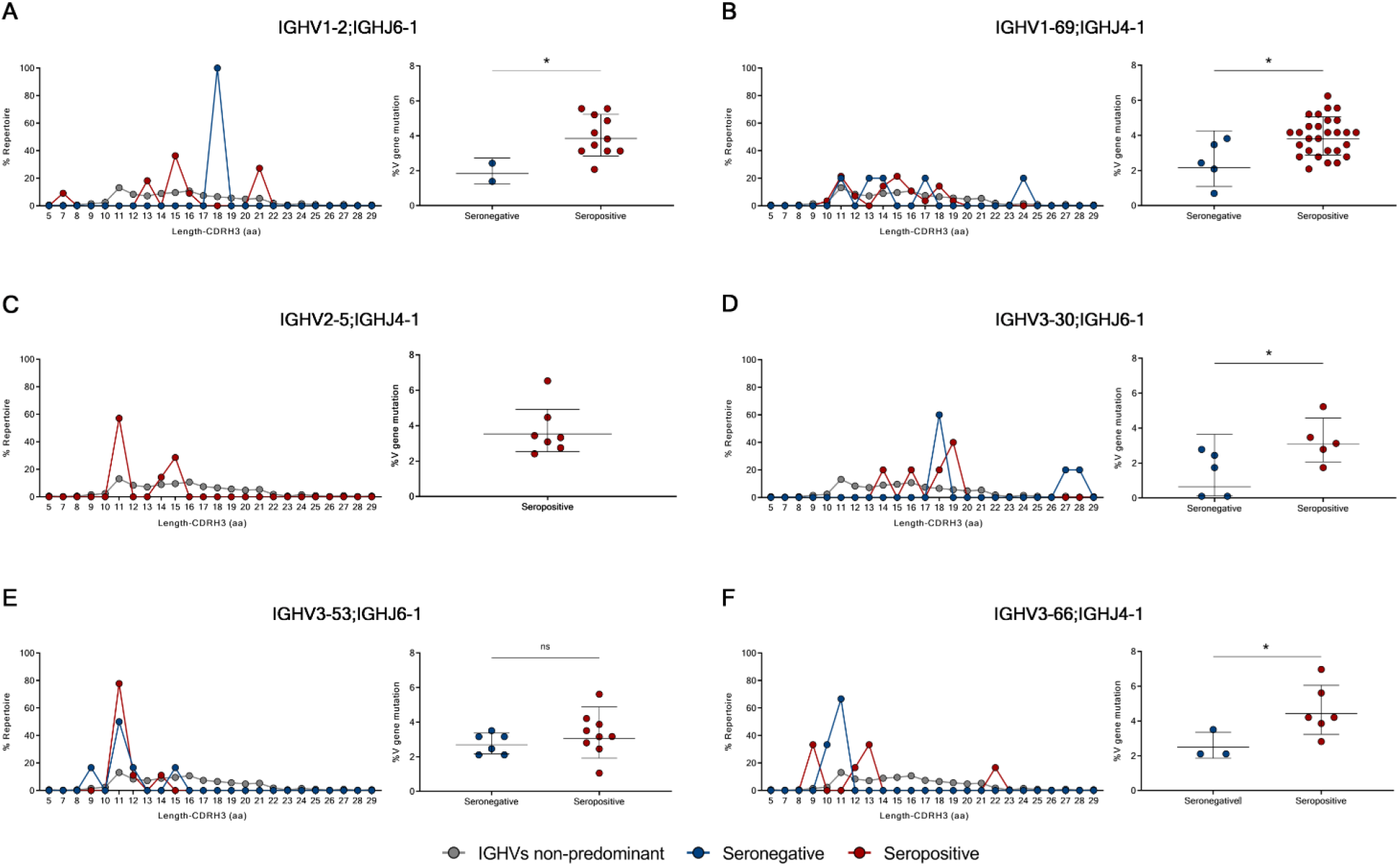
Heavy chain CDR3 length and somatic hypermutation levels of predominant gene derived nAbs. (A - F) Graphs show the amino acidic heavy chain CDR3 length (left panel) and the somatic hypermutation level (right panel) of nAbs derived from the IGHV1-2;IGHJ6-1 (A), IGHV1-69;IGHJ4-1 (B), IGHV2-5;IGHJ4-1 (C), IGHV3-30;IGHJ6-1 (D), IGHV3-53;IGHJ6-1 (E) and IGHV3-66;IGHJ4-1 (F) gene families.

**Figure S6.**
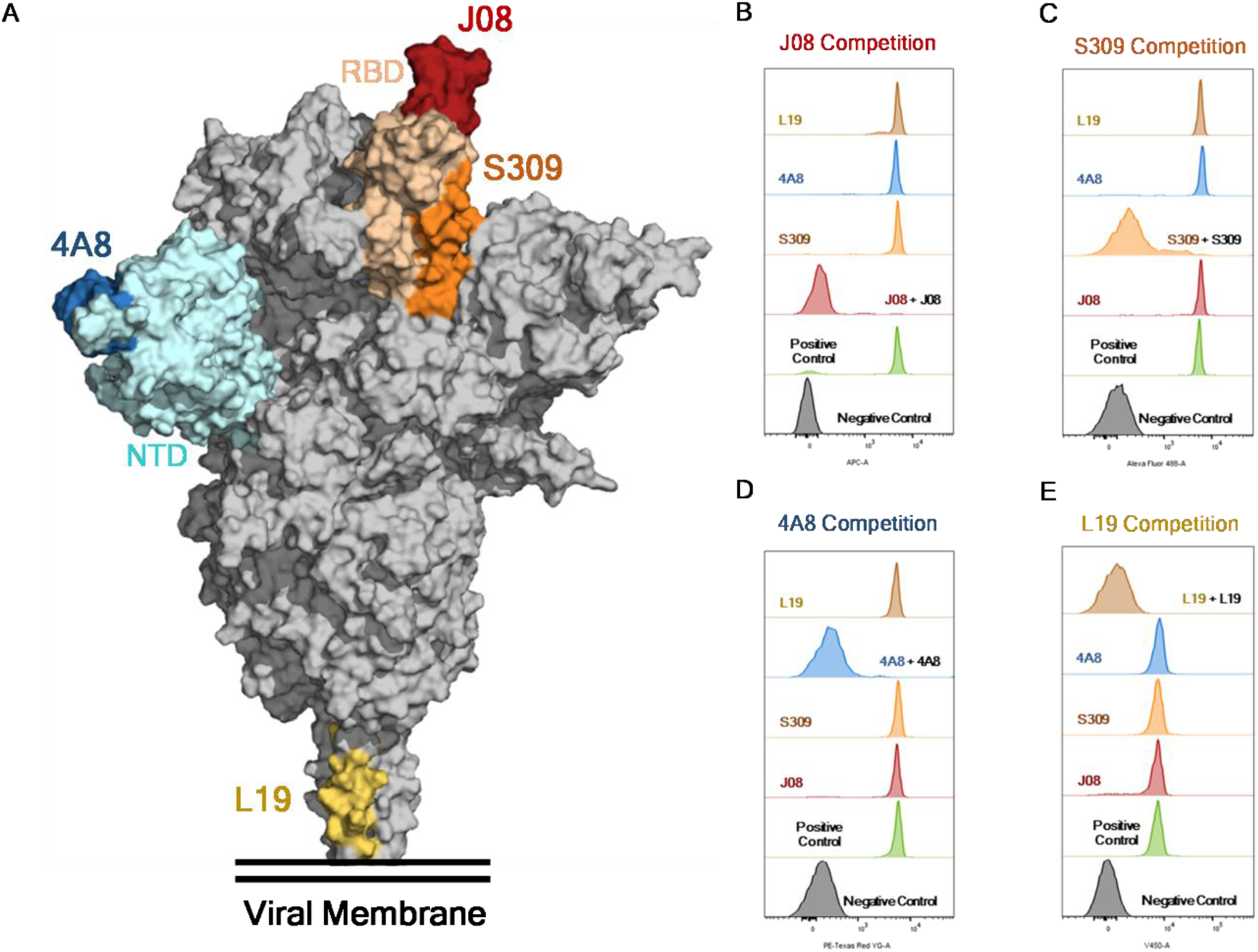
Heavy chain CDR3 length and somatic hypermutation levels of predominant gene derived nAbs. (A) Schematic representation of the epitopes recognized by J08 (dark red), S309 (orange), 4A8 (dark blue) and L19 (gold), mAbs on the S protein surface. (B - E) Representative cytometer peaks per each of the four mAbs used for the competition assay. Positive (beads conjugated with only primary labeled antibody) and negative (un-conjugated beads) controls are shown as green and gray peaks, respectively.

## REFERENCES

1 Wajnberg, A. et al. Robust neutralizing antibodies to SARS-CoV-2 infection persist for months. 370, 1227–1230, doi:10.1126/science.abd7728 %J Science (2020).

2 Xiaojie, S., Yu, L., lei, Y., Guang, Y. & Min, Q. Neutralizing antibodies targeting SARS-CoV-2 spike protein. Stem Cell Research 50, 102125, doi:https://doi.org/10.1016/j.scr.2020.102125 (2021).

3 Andreano, E. et al. Extremely potent human monoclonal antibodies from COVID-19 convalescent patients. Cell 184, 1821–1835.e1816, doi:https://doi.org/10.1016/j.cell.2021.02.035 (2021).

4 Yang, Y. & Du, L. SARS-CoV-2 spike protein: a key target for eliciting persistent neutralizing antibodies. Signal Transduction and Targeted Therapy 6, 95, doi:10.1038/s41392-021-00523-5 (2021).

5 Clausen, T. M. et al. SARS-CoV-2 Infection Depends on Cellular Heparan Sulfate and ACE2. Cell 183, 1043–1057.e1015, doi:https://doi.org/10.1016/j.cell.2020.09.033 (2020).

6 Wrapp, D. et al. Cryo-EM structure of the 2019-nCoV spike in the prefusion conformation. 367, 1260–1263, doi:10.1126/science.abb2507 %J Science (2020).

7 Piccoli, L. et al. Mapping Neutralizing and Immunodominant Sites on the SARS-CoV-2 Spike Receptor-Binding Domain by Structure-Guided High-Resolution Serology. Cell 183, 1024–1042.e1021, doi:10.1016/j.cell.2020.09.037 (2020).

8 Tortorici, M. A. et al. Broad sarbecovirus neutralization by a human monoclonal antibody. Nature, doi:10.1038/s41586-021-03817-4 (2021).

9 Andreano, E. & Rappuoli, R. Immunodominant antibody germlines in COVID-19Immunodominant antibody germlines in COVID-19. Journal of Experimental Medicine 218, doi:10.1084/jem.20210281 (2021).

10 Yuan, M. et al. Structural basis of a shared antibody response to SARS-CoV-2. 369, 1119– 1123, doi:10.1126/science.abd2321 %J Science (2020).

11 Callaway, E. Coronavirus variants get Greek names - but will scientists use them? Nature 594, 162, doi:10.1038/d41586-021-01483-0 (2021).

12 Stamatatos, L. et al. mRNA vaccination boosts cross-variant neutralizing antibodies elicited by SARS-CoV-2 infection. 372, 1413–1418, doi:10.1126/science.abg9175 %J Science (2021).

13 Goel, R. R. et al. Distinct antibody and memory B cell responses in SARS-CoV-2 naïve and recovered individuals after mRNA vaccination. 6, eabi6950, doi:10.1126/sciimmunol.abi6950 %J Science Immunology (2021).

14 Urbanowicz, R. A. et al. Two doses of the SARS-CoV-2 BNT162b2 vaccine enhances antibody responses to variants in individuals with prior SARS-CoV-2 infection. eabj0847, doi:10.1126/scitranslmed.abj0847 %J Science Translational Medicine (2021).

15 Crotty, S. Hybrid immunity. 372, 1392–1393, doi:10.1126/science.abj2258 %J Science (2021).

16 Wang, Z. et al. Naturally enhanced neutralizing breadth against SARS-CoV-2 one year after infection. Nature 595, 426–431, doi:10.1038/s41586-021-03696-9 (2021).

17 Zhang, Q. et al. Potent and protective IGHV3-53/3-66 public antibodies and their shared escape mutant on the spike of SARS-CoV-2. Nature Communications 12, 4210, doi:10.1038/s41467-021-24514-w (2021).

18 Pinto, D. et al. Cross-neutralization of SARS-CoV-2 by a human monoclonal SARS-CoV antibody. Nature 583, 290–295, doi:10.1038/s41586-020-2349-y (2020).

19 Chi, X. et al. A neutralizing human antibody binds to the N-terminal domain of the Spike protein of SARS-CoV-2. 369, 650–655, doi:10.1126/science.abc6952 %J Science (2020).

20 Voss, W. N. et al. Prevalent, protective, and convergent IgG recognition of SARS-CoV-2 non-RBD spike epitopes. 372, 1108–1112, doi:10.1126/science.abg5268 %J Science (2021).

21 Li, B. et al. Viral infection and transmission in a large, well-traced outbreak caused by the SARS-CoV-2 Delta variant. 2021.2007.2007.21260122, doi:10.1101/2021.07.07.21260122%JmedRxiv (2021).

22 Meng, B. et al. Recurrent emergence of SARS-CoV-2 spike deletion H69/V70 and its role in the Alpha variant B.1.1.7. Cell Reports 35, doi:10.1016/j.celrep.2021.109292 (2021).

23 Steichen, J. M. et al. A generalized HIV vaccine design strategy for priming of broadly neutralizing antibody responses. 366, eaax4380, doi:10.1126/science.aax4380 %J Science (2019).

24 Havenar-Daughton, C., Abbott, R. K., Schief, W. R. & Crotty, S. When designing vaccines, consider the starting material: the human B cell repertoire. Current opinion in immunology 53, 209–216, doi:10.1016/j.coi.2018.08.002 (2018).

25 Huang, J. et al. Isolation of human monoclonal antibodies from peripheral blood B cells. Nature Protocols 8, 1907–1915, doi:10.1038/nprot.2013.117 (2013).

26 Planas, D. et al. Sensitivity of infectious SARS-CoV-2 B.1.1.7 and B.1.351 variants to neutralizing antibodies. Nature Medicine 27, 917–924, doi:10.1038/s41591-021-01318-5 (2021).

27 Tiller, T. et al. Efficient generation of monoclonal antibodies from single human B cells by single cell RT-PCR and expression vector cloning. J Immunol Methods 329, 112–124, doi:10.1016/j.jim.2007.09.017 (2008).

28 Wardemann, H. & Busse, C. E. Expression Cloning of Antibodies from Single Human B Cells. Methods in molecular biology (Clifton, N.J.) 1956, 105–125, doi:10.1007/978-1-4939-9151-8_5 (2019).

29 Kepler, T. B. Reconstructing a B-cell clonal lineage. I. Statistical inference of unobserved ancestors. F1000Res 2, 103–103, doi:10.12688/f1000research.2-103.v1 (2013).

30 Kepler, T. B. et al. Reconstructing a B-Cell Clonal Lineage. II. Mutation, Selection, and Affinity Maturation. 5, doi:10.3389/fimmu.2014.00170 (2014).

